# Cataract among elderly in Sri Lanka

**DOI:** 10.1101/540518

**Authors:** A.A. Nilanga Nishad, S.A. Hewage, K. Arulmoly, M.S. Amaratunge, J de Silva, KTAA Kasturiratne, PK Abeysundara, AR Wickramasinghe

## Abstract

Out of 39 billion people who are blind around the world, 20 billion (51.3%) is due to cataract, which is preventable. This study intended to assess the prevalence and factors associated with cataract among elderly in a divisional secretariat area in Sri Lanka. This community based cross sectional study assessed randomly selected470 adults over 60 years of age. Diagnosis of cataract was made by a slit lamp examination by medical officers, and classified according to Oxford Lens Opacity Classification system (LOCS III). Majority was between 60-69 age groups and 71% was females. The prevalence of cataract was estimated to be 80.6% including operated eye and 73.6% excluding the operated eye, with a female preponderance in lower age categories. Commonest type of cataract was the nuclear type (n=422; 44.9%), with a majority in grade 2 (218; 23.2%). The prevalence of cataract surgery in the diseased population was as low as 7%. Cataract leading to blindness is very prevalent among adults over 60 years of age in the studied area. Females tend to develop the disease at an early age than males. These findings warrant screening programme for elderly at community level, targeting females at a younger age than males. Future studies are recommended to assess the coverage and barriers for cataract surgeries at national level, which would be immensely useful in planning and improving health services.

## Introduction

Despite efforts taken by many national and international agencies, blindness scales up to be a major global health issue. Approximately 39 million peoplesuffer from blindness worldwide, and preventable causes are as high as 80% of this global burden ^[1]^. Blindness or poor visual functions leads to many adverse outcomes including limitations in mobility, activities of daily living, physical function and poor quality in life. The World Health Organization (WHO) estimates cataract to be responsible for 51% of world blindness ^[2]^, which represents 20 million people around the globe. This makes cataract, defined as a clouding or loss of transparency of the lens in the eye as a result of tissue breakdown and protein clumping ^[3]^, the leading preventable cause of blindness. Cataract itself can be due to many risk factors, but ageing remains the number one non-modifiable risk factor. Ever growing rate of the elderly population in the world challenges health systems around the globe in planning preventive strategies and implementing them to tackle the problem of blindness due to cataract effectively. In its book on ‘Strategies for the prevention of blindness in national programmes’, the WHO has identified national programmes based on a primary health care approach as the best strategy for reaching the many people who can benefit from simple, inexpensive and effective measures delivered by trained health workers ^[4]^. Sri Lanka has a world renowned public health system with a network of grass root level workers delivering health services to people at community level^[5]^,but blindness has never beenrecognized as a priority area for the health care workers. This is irrespective of the facts that Sri Lanka records the fastest ageing rate in South East Asia, and expects one in every four citizens to be over 60 years of age by 2025 ^[6]^. There are no established, routine screening programmes or public awareness campaigns conducted by the public health system at community levels to address cataract.

Cataract remains to be the leading cause for blindness in Low-Middle Income Countries (LMICs) too ^[7]^. Priority blinding diseases in Sri Lanka are known to be cataract, blindness in children, glaucoma, diabetic retinopathy and refractive errors ^[8]^. However, Sri Lanka does not have national level data for the prevalence of cataract. Few individual studies report a range of prevalence rates as high as56% ^[9]^ to 33.1%^[10]^for cataract among elderly, and the last study was reported in 2009. Understanding the epidemiological profile of a disease is the crucial foundation in planning effective preventive strategies. Therefore, assessing the epidemiology of cataract among elderly in Sri Lanka is a timely need to target preventive measures. This study assessed the prevalence and distribution of cataract among the population over 60 years of age in a divisional secretariat area.

## Methods and materials

A cross sectional study was carried out in Mahara Divisional Secretariat (DS) division in Gampaha district, one of the district with highest population in Sri Lanka, with an elderly population of 290,386 ^[11]^. Multi stage cluster sampling method was used to draw the study population from the target population of adults over 60 years of age living in Gampaha district. Critically ill, bed bound and mentally unsound elders were excluded from the sampling frame. Administratively, DS divisions are further divided into smaller Grama Niladhari (GN) divisions. At the first stage, 18 such GN divisions were selected out of 92 GN divisions in Mahara DS division by simple random sampling method using computer generated random numbers without replacement. At the second stage, 30 participants from each GN division were randomly selected using the voters’ list as the sampling frame, without considering the size of elderly population in each GN division. Calculated sample size of 470 was achieved after inviting 540 elders.

Visual acuity was measured and slit lamp examination was carried out by medical officers with an experience in ophthalmology for more than 5 years on all study participants. Dilation of the eye was carried out before examination. Randomly chosen subsets of samples were examined by a consultant ophthalmologist as a quality assurance process. Diagnosis of cataracts was classified as nuclear, cortical or sub-capsular cataract and was graded according to Oxford Lens Opacity Classification system (LOCS III)^[12]^.

An Interviewer administered, pre-tested questionnaire was used to collect data. Content validity of the questionnaire was ensured by a thorough literature search, taking into consideration opinions of specialist Ophthalmologists in the development of the instruments. Data collection was carried out during a period of 10 weeks starting from 1st August 2012. All possible information were cross checked with written records, for example, the age with participants’ national identity cards, diagnoses of diseases with medical records and previous cataract surgeries with diagnosis cards, to preserve the validity of data.

Descriptive data were presented as frequencies and percentages. Continuous data were summarized using mean with standard deviations and range. Significance of association was tested using the Chi square (χ2) test for categorical variables and Student’s t-test for continuous variables. The p value was considered as significant at 5% level.

Participants who were tested positive for cataract were counseled for surgery and were given a date for surgery (free of charge inclusive of the lens). Participants with other visual problems like diabetic retinopathy, age related macular degeneration and glaucoma were referred to necessary specialist units at the Seedwa Vijayakumaratunge Memorial Hospital for further management. Spectacles were distributed to participants who required them, free of charge. Ethical clearance was obtained from the Ethical Review Committee of Faculty of Medicine, University of Kelaniya, Sri Lanka (reference number P108/07/2012).

## Results

A total of 540 people were invited to participate in the study, of which 470 people participated resulting in a response rate of 87%. Table 1 display there was no difference between respondents and non-respondents with regard to the basic demographic characteristics.

**Table 1.**
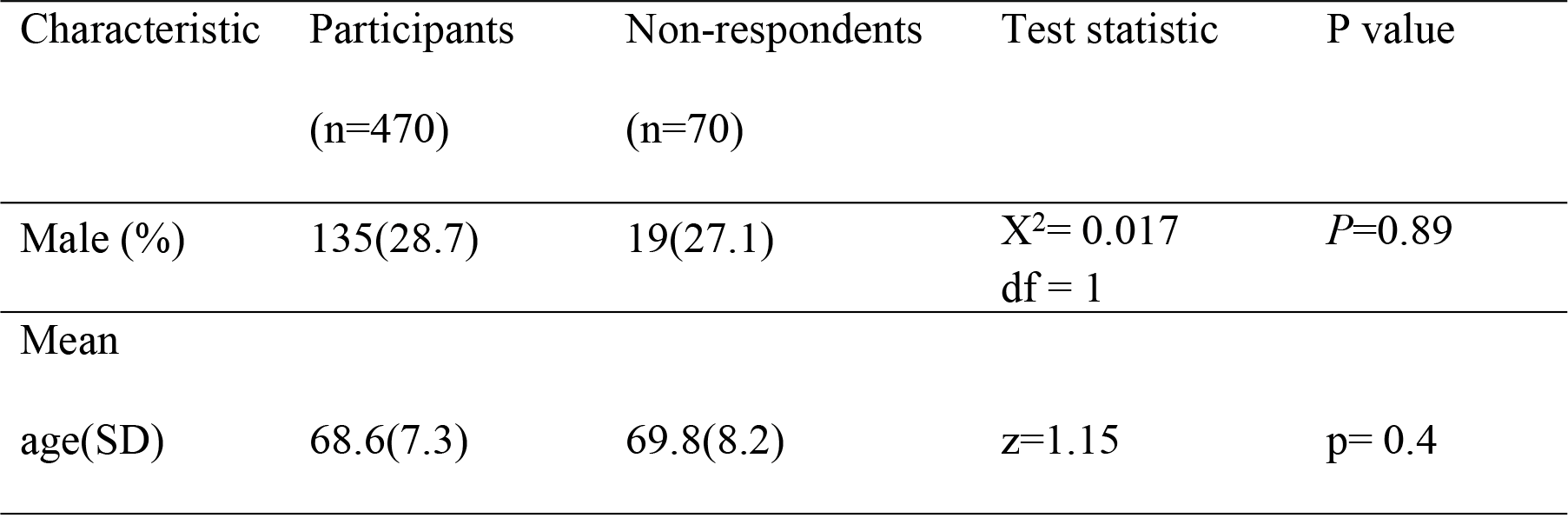
Comparison of study participants and non-respondents by age and gender.

Majority of participants were between 60-69 years of age, and 71% of the sample was females (Table 2).

**Table 2.**
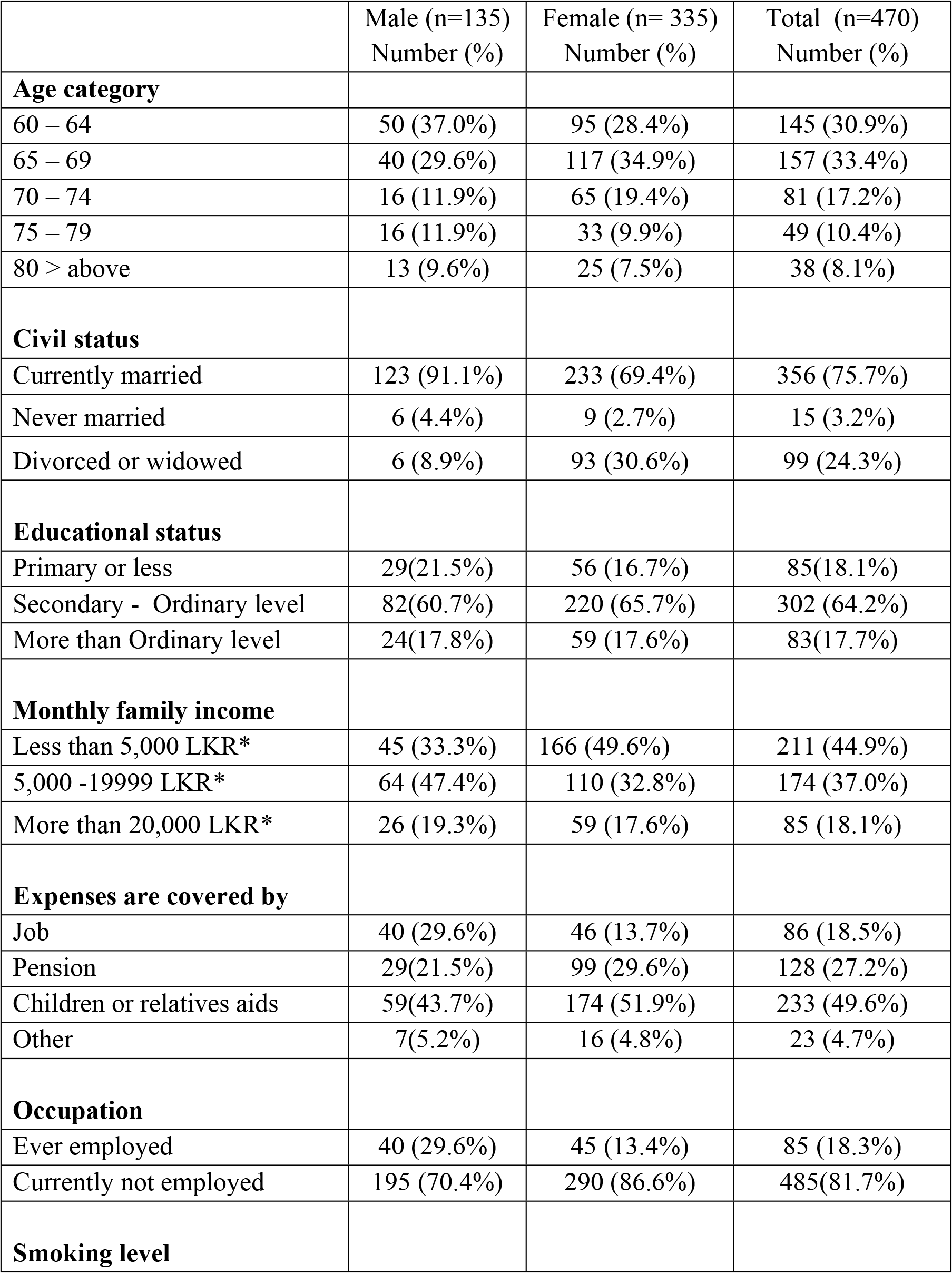
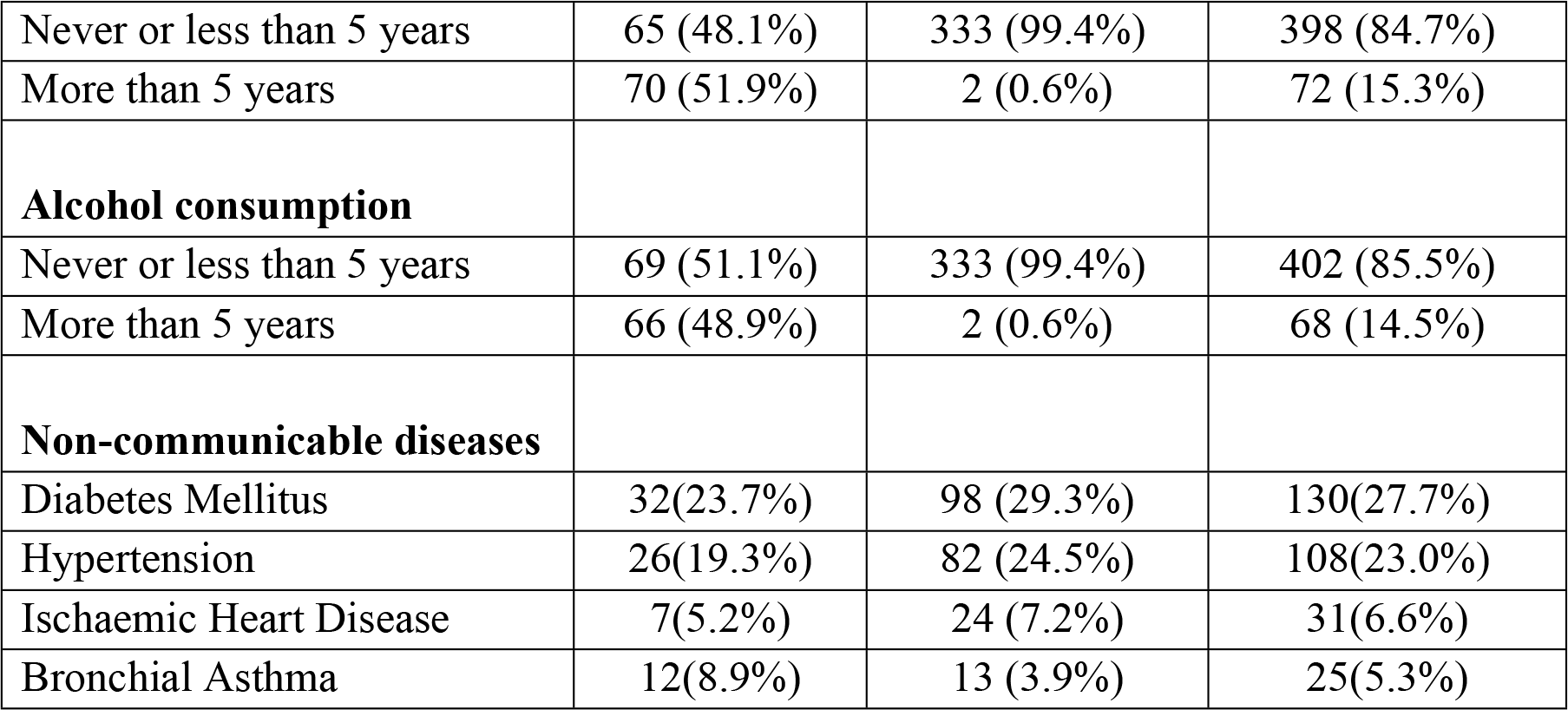
Socio-demographic profile of the participants by gender.

|  | Male (n=135)<br>Number (%) | Female (n= 335)<br>Number (%) | Total (n=470)<br>Number (%) |
| --- | --- | --- | --- |
| **Age category** |  |  |  |
| 60 – 64 | 50 (37.0%) | 95 (28.4%) | 145 (30.9%) |
| 65 – 69 | 40 (29.6%) | 117 (34.9%) | 157 (33.4%) |
| 70 – 74 | 16 (11.9%) | 65 (19.4%) | 81 (17.2%) |
| 75 – 79 | 16 (11.9%) | 33 (9.9%) | 49 (10.4%) |
| 80 > above | 13 (9.6%) | 25 (7.5%) | 38 (8.1%) |
| **Civil status** |  |  |  |
| Currently married | 123 (91.1%) | 233 (69.4%) | 356 (75.7%) |
| Never married | 6 (4.4%) | 9 (2.7%) | 15 (3.2%) |
| Divorced or widowed | 6 (8.9%) | 93 (30.6%) | 99 (24.3%) |
| **Educational status** |  |  |  |
| Primary or less | 29(21.5%) | 56 (16.7%) | 85(18.1%) |
| Secondary - Ordinary level | 82(60.7%) | 220 (65.7%) | 302 (64.2%) |
| More than Ordinary level | 24(17.8%) | 59 (17.6%) | 83(17.7%) |
| **Monthly family income** |  |  |  |
| Less than 5,000 LKR* | 45 (33.3%) | 166 (49.6%) | 211 (44.9%) |
| 5,000 -19999 LKR* | 64 (47.4%) | 110 (32.8%) | 174 (37.0%) |
| More than 20,000 LKR* | 26 (19.3%) | 59 (17.6%) | 85 (18.1%) |
| **Expenses are covered by** |  |  |  |
| Job | 40 (29.6%) | 46 (13.7%) | 86 (18.5%) |
| Pension | 29(21.5%) | 99 (29.6%) | 128 (27.2%) |
| Children or relatives aids | 59(43.7%) | 174 (51.9%) | 233 (49.6%) |
| Other | 7(5.2%) | 16 (4.8%) | 23 (4.7%) |
| **Occupation** |  |  |  |
| Ever employed | 40 (29.6%) | 45 (13.4%) | 85 (18.3%) |
| Currently not employed | 195 (70.4%) | 290 (86.6%) | 485(81.7%) |
| **Smoking level** |  |  |  |
| Never or less than 5 years | 65 (48.1%) | 333 (99.4%) | 398 (84.7%) |
| More than 5 years | 70 (51.9%) | 2 (0.6%) | 72 (15.3%) |
| **Alcohol consumption** |  |  |  |
| Never or less than 5 years | 69 (51.1%) | 333 (99.4%) | 402 (85.5%) |
| More than 5 years | 66 (48.9%) | 2 (0.6%) | 68 (14.5%) |
| **Non-communicable diseases** |  |  |  |
| Diabetes Mellitus | 32(23.7%) | 98 (29.3%) | 130(27.7%) |
| Hypertension | 26(19.3%) | 82 (24.5%) | 108(23.0%) |
| Ischaemic Heart Disease | 7(5.2%) | 24 (7.2%) | 31(6.6%) |
| Bronchial Asthma | 12(8.9%) | 13 (3.9%) | 25(5.3%) |

Prevalence of any form of cataract in at least one eye including operated eyes was 80.6% (95%CI 76.8-84%) and was 73.6% (95%CI 69.5-77.4%) when operated eye was excluded. Nuclear cataract was the commonest single type of cataract with a prevalence rate of 44.9% (Table 3).

**Table 3. Prevalence of different sub-types of cataract by gender among the participants**

As expected, the prevalence of cataract increased with increasing age (table 4). The prevalence among females was higher in lower age groups than males, and the pattern reverses in older age groups. Table 4 describes the distribution of different types of cataract across different age groups among males and females separately. It also shows the very low coverage of cataract surgeries among the affected population.

**Table 4. Prevalence of different types of cataracts by age and gender**

## Discussion and conclusion

The response rate was 87% and there was no difference between the non-participants and study participants in terms of age and sex. The mean age (SD) of the study participants was 68.6 (7.3) years Eighty percent of participants were below 74 years of age. The prevalence of cataract was estimated to be 80.6% including operated eye and 73.6% excluding the operated eye (Table 2), with a female preponderance in lower age categories. The commonest type of cataract was the nuclear type cataract followed by the mixed type and sub-capsular type (Table 3). The prevalence of cataract surgery in the diseased population was 7%. Nuclear-cortical, nuclear-sub-capsular and all three subtypes together were present but the cortical-sub-capsular combination was not seen together. As expected, the prevalence of cataract increased with increasing age and grades 2 and 3 were the common grades of cataract seen in this study. Exclusion of bed bound elders and those with critical illnesses might have diluted the prevalence of cataract among the elderly.

The prevalence of cataract was estimated to be 80.6% including operated eye and 73.6% excluding the operated eye (Table 4). This is the highest reported prevalence of cataract in a study conducted in Sri Lanka. Previous reported prevalence rates varied between 56% ^[9]^ to 33.1% ^[10]^. This could be due to differences in methods of studies. For example, the study by Edussuriya et al (2009) which reports a lower prevalence rate for cataract of 33.1% included people above 40 years of age, which must have diluted the results as the disease is associated with ageing process. Moreover, different methods used to diagnose cataract clinically must have affected prevalence rates. This study was conducted among people above 60 years of age, the cutoff age to define elderly by the WHO, and cataract was diagnosed by slit lamp examination, which is the gold standard method to clinically diagnose the condition.

The commonest type of cataract identified by this study was the nuclear type cataract followed by the mixed type and sub-capsular type. Females had a higher prevalence of cataract (Table 4). Nuclear-cortical, nuclear-sub-capsular and all three subtypes together were present but the cortical-sub-capsular combination was not seen together (Table 3). Grades 2 and 3 were the common grades of cataract seen in this study. As expected, the prevalence of cataract increased with increasing age (Table 4). Since we have excluded elders with critical illnesses and those who were bed bound, who are more likely to be in the higher end of the age range, we may have underestimated the prevalence of cataracts in the older age group. As expected, prevalence of cataract was positively associated with increasing age (Table 4). In their population based study, Xu et al.reported acataract prevalence of 97.8% in people over 70 years of age, the highest reported estimate as of date ^[17]^. A similar pattern was seen in other Sri Lankan studies as well even though the prevalence was not as high as that reported by Xu et al.^[7,8,9]^. As Table 4 display, the overall prevalence of cataract in both sexes was similar. Hapugodaalso reported equal prevalence of cataract in both sexes in Sri Lanka in 2009^[18]^, but Nanayakkarareported a female preponderance in cataract prevalence over males (65%:48.2%) in 2004^[9]^. Praveen et al. also reported a higher prevalence of cataract among females in India^[15]^. The prevalence of cataract was found to be higher among the females in younger age groups in this study. Further studying this factor will assess whether there is a need to start screening for cataract in females at an earlier age than males.

The very low coverage of cataract surgeries among the study population is noteworthy. Only 7.4% of males diagnosed with cataract had undergone a surgery, while the same for females was 6.9%. This finding definitely opens doors to further research to look into determinants of this poor coverage, which could be used to plan and improve the services provided to patients in the future.

In this study, nuclear cataract was the commonest single type of cataract (44.9%) followed by the mixed type (27%). Nuclear cataract was reported as the most common sub type in most of the regional studies as well. An Indian study describes a prevalence of nuclear cataract and mixed type to be 48% and 38% respectively^[15]^ and in Chinaa prevalence of nuclear cataract of 50% was reported in 2006 ^[17]^. Taiwan reported a nuclear cataract prevalence of 38%^[16]^. However, Husain et al. reported the mixed type of cataract to be the commonest type of cataract in Indonesia ^[14]^. Previous Sri Lankan studies have not described the type of cataracts detected. The prevalences of the different types of cataracts and the cataract surgery rate were similar in the two sexes according to the current study (Table 4).

## Strength and limitations

A major strength of this study is that it used the gold standard test for diagnosis of cataract which is the detailed clinical examination including ophthalmoscopy and slit lamp examination by a clinician. Many previous studies conducted in Sri Lanka used a simpler methodology to screen for cataract. Inclusion of adults over 60 years of age also clearly defines the elderly population according to the WHO criteria. On the other hand, the external validity is limited as the geographical area of the study was confined to a DS division. However, capturing the study population from a wider geographical area was not possible owing to the time and financial constraints. In addition, aetiology of cataract is not known to be related to any environmental or regional factors so far. Therefore, this limitation is not a serious threat to the generalizability.

## Conclusions and recommendations

Cataractleading to blindness is very prevalent among adults over 60 years of age in the studied area, and the rate increases with ageing. There is no difference in the prevalence of overall cataract in both sexes, but females tend to suffer from the disease at an early stage than males. Since the condition can be screened using very simple tests and curative surgeries can be offeredfor diagnosed patients at majority of hospitals in Sri Lanka, this high prevalence justifies initiating a screening programme for elderly at community level, targeting females at a younger age than males.

The coverage of curative surgeries for cataract among the studied population is very low. Further studies are recommended to assess the situation throughout Sri Lanka and also to assess the determinants which would be immensely useful in planning and improving health services to the population.

## Acknowledgement

The authors wish to acknowledge the support received from the medical staff at the Seeduwa Vijayakumaratune Memorial Hospital, Sri Lanka, including consultant Ophthalmologists, for their participation at the study for eye examinations and follow up of referred patients, and the director of the Hospital for his administrative permissions to use required equipments.

